# Establishing an *ex vivo* platform for molecular profiling of skin structural cells using frozen human skin biopsies

**DOI:** 10.64898/2026.10.30.763216

**Authors:** Ayako Takamori, Hannah DeBerg, Peter A. Morawski, Mitch Fahning, James D. Schlenker, William P. Schmitt, Iris Gratz, Daniel Campbell

## Abstract

Skin structural cells, including keratinocytes, melanocytes, endothelial cells, and fibroblasts, play critical roles in the development of chronic inflammatory skin disease. Inflammatory stimuli induce transcriptional reprogramming and epigenetic changes in skin structural cells, which contribute to disease pathogenesis. Although two-dimensional (2D) and three-dimensional (3D) skin culture models have been widely used to investigate inflammatory responses in skin structural cells, *ex vivo* systems that maintain skin structure and cellular composition remain limited. To address this gap, we explored using frozen skin biopsies as an experimental platform to study the responses of skin structural cells to inflammatory stimuli. Using a simple air-liquid interface culture method on permeable Transwell inserts, we showed that viable skin structural cells that retain cell-type-specific transcriptional profiles can be recovered from frozen skin biopsies after prolonged *in vitro* culture. Stimulation with IFN-γ induced robust transcriptional and chromatin accessibility changes that were largely resolved after stimulus removal, with transcriptional responses to secondary stimulation comparable to the primary stimulation. Treatment of cultured skin biopsies with histone-modifying enzyme inhibitors effectively altered global histone modification levels, but did not substantially affect the transcriptional resolution of IFN-γ-induced responses. Collectively, these findings establish cryopreserved human skin biopsies as a scalable *ex vivo* platform that enables mechanistic studies of inflammatory responses of multiple populations of skin structural cells.

## Introduction

The skin acts as a physical and immunological barrier against the external environment. It consists of multiple cell types, including structural cells, such as keratinocytes, fibroblasts, melanocytes, and endothelial cells, as well as immune cells. In addition to providing structural support, skin structural cells actively communicate with each other and with local immune cells to maintain tissue homeostasis and regulate inflammatory responses and tissue repair. [1, 2]. Increasing evidence indicates that skin structural cells contribute to the pathogenesis of chronic inflammatory skin diseases, including psoriasis and atopic dermatitis [3, 4]. Furthermore, these cells also express immune-related gene programs that enable them to sense and respond to inflammatory stimuli [5].

The pathogenesis of skin inflammatory disease is driven by inflammatory cytokines released by immune and skin structural cells as well as by pathogen-associated molecular patterns (PAMPs), which are recognized by pattern recognition receptors such as Toll-like receptors (TLRs) that are expressed by immune and structural cells [6, 7]. Chronic inflammatory skin diseases are associated with transcriptional reprogramming of skin structural cells, including keratinocytes and fibroblasts [8, 9]. Beyond transcriptional changes, changes in chromatin accessibility and histone modifications have been identified in structural cells in chronic inflammatory skin disease [10, 11]. In line with these observations, cytokine stimulation of cultured human keratinocytes induces transcriptional and epigenetic alterations that partially recapitulate those observed in psoriatic lesions [12]. In addition, studies in mice have demonstrated that inflammatory stimuli induce persistent chromatin accessibility changes and alter responses to subsequent stimulation in epidermal stem cells and fibroblasts, a phenomenon referred to as inflammatory memory [13–15]. Collectively, these observations suggest that inflammatory stimuli induce transcriptional and epigenetic alterations in skin structural cells that initiate and amplify development of inflammatory skin diseases. The clinical success of targeting cytokine-driven inflammatory circuits highlights their importance in the pathogenesis of inflammatory skin diseases[16, 17]. However, experimental systems that enable evaluation of cytokine-driven transcriptional and epigenetic changes in physiologically relevant human skin structural cells remain limited.

Several experimental systems are used to evaluate inflammatory stimulus-induced responses in skin structural cells, including two-dimensional (2D) skin culture systems, three-dimensional (3D) engineered culture systems, and skin biopsy systems [18]. Each has advantages and limitations. 2D skin culture systems are relatively easy to handle, and suitable for initial high-throughput analysis, but lack a structural context and may use immortalized cell lines [19, 20]. 3D-engineered culture systems are more physiologically relevant compared to 2D skin culture systems, but remain limited in cellular diversity [20]. In contrast, skin biopsies maintain more physiologically relevant tissue organization together with diverse resident cells, including melanocytes and endothelial cells which are often absent from conventional 2D or 3D skin culture systems [21], but these can be limited by tissue availability. Therefore, experimental systems that combine physiological relevance with improved experimental flexibility, throughput, and reproducibility are needed. Frozen skin biopsies may provide such an *ex vivo* platform by enabling sample storage while preserving key features of human skin tissue. To address this need, we established an *ex vivo* system using frozen human skin biopsies from healthy donors to evaluate inflammatory stimulus-induced transcriptional and epigenetic changes in multiple populations of skin structural cells by bulk RNA-seq and ATAC-seq.

## Materials and Methods

### Human skin samples

Human skin samples were obtained from healthy donors undergoing elective panniculectomy or abdominoplasty. All samples were obtained with written informed consent at Virginia Mason Franciscan Health (Seattle, WA). All study protocols were conducted in accordance with the Declaration of Helsinki principles and were approved by the Institutional Review Board of Benaroya Research Institute (Seattle, WA, USA).

### Skin punch biopsies

Fresh surgical discard abdominal skin samples were treated with PBS containing 0.1% Primocin (Invitrogen; ant-pm-1) for 5 minutes to eliminate contaminating microorganisms. Tissue processing was performed on ice, and skin was periodically moistened with 1x PBS to prevent drying. After removal of subcutaneous fat using scissors, a 4-mm biopsy punch (MedBlades) was used to obtain tissue biopsies from each sample Twelve 4-mm skin punch biopsies were cryopreserved in each cryovial in bovine calf serum (BCS) containing 7% DMSO and stored in liquid nitrogen until use. Cryopreserved biopsies were rapidly thawed in a 37 °C water bath and washed twice with PBS prior to further processing.

### Culture conditions

Skin biopsies were either processed immediately after thawing (day 0) or cultured for up to day 14 at 37 °C in 5% CO_2_. For culture, 700 µL of Green’s Plus medium (DMEM and Ham’s F-12 supplemented with 10% fetal bovine serum (FBS), L-glutamine, sodium pyruvate, adenine, hydrocortisone, insulin, cholera toxin, triiodothyronine (T3), penicillin-streptomycin, and 10 ng/mL human epidermal growth factor (hEGF) [22] was added to the lower chamber of 12 well Transwell plates (Corning). Skin punch biopsies were placed on 3.0 µm pore size Transwell inserts, with the dermal side down and the epidermis exposed to air. The medium was changed every other day.

### IFN-γ stimulation, removal, and secondary stimulation

For stimulation, 100 ng/mL recombinant human IFN-γ (BioLegend) was added to the lower chamber 24 hours prior to analysis. For IFN-γ removal experiments, skin biopsies were stimulated with IFN-γ on day 4 for 24 hours, briefly rinsed on the bottom surface with PBS to remove IFN-γ, transferred to new Transwell inserts, and cultured for an additional 5 days, with medium changed every other day. On day 10, the biopsies were harvested for downstream analysis. For secondary stimulation experiments, biopsies were stimulated with IFN-γ on day 4 for 24 hours, followed by washing and cultured as described above. On day 10, the biopsies were restimulated with IFN-γ for 24 hours, then digested to generate single cell suspensions for downstream analysis.

### Histone-modifying enzyme inhibitor treatment

For histone-modifying enzyme inhibitor treatment, skin biopsies were stimulated with IFN-γ on day 4 for 24 hours. On day 5, the biopsies were washed with PBS and transferred to new Transwell inserts. Immediately after transfer, the HDAC inhibitor entinostat (MedChemExpress) (1 µM) or the EZH2 inhibitor tazemetostat (MedChemExpress) (0.1 µM) or DMSO as a vehicle control was added to the medium in the lower chamber. The medium containing either DMSO or the indicated inhibitor was replaced every other day on days 5, 7, and 9. On day 10, biopsies were then digested and the epidermal cell suspensions were either analyzed by flow cytometry to assess histone modification levels in gated keratinocytes, or keratinocytes were sorted for bulk RNA-seq.

### Isolation of skin structural cells from skin biopsies

Skin biopsies were digested for 90 minutes at 37 °C with 25 U/mL dispase II (Gibco) in calcium– and magnesium free HBSS and the epidermis was separated from the dermis. For isolation of keratinocytes and melanocytes, epidermal sheets were further digested with either TrypLE Express (Gibco) for 15 minutes at 37 °C with shaking at 150 rpm for bulk RNA-seq, or 0.25% trypsin-EDTA (Gibco) for 3 minutes at 37 °C without shaking for bulk ATAC-seq. After trypsin inactivation in medium containing 10% FBS, cells were filtered through a 70 µm filter, followed by sorting or downstream processing. For isolation of endothelial cells and fibroblasts, dermal tissue was digested with 2.2 mg/mL collagenase (Worthington) and 50 µg/mL DNase I (Sigma-Aldrich) in RPMI containing 5% human serum for 90 minutes at 37 °C with shaking at 150 rpm. The tissue was then mechanically dissociated by further mincing with scissors and filtered through a 100 µm filter. Cells were washed with medium containing 3% BCS and collected for sorting.

### Flow cytometry and cell sorting

Following isolation, cells were incubated with a fixable viability dye (Fixable Viability Dye eFluor 780, Invitrogen) and an Fc receptor blocking reagent (Human TruStain FcX, BioLegend) for 10 minutes at room temperature. Without washing, cells were then stained with antibodies at 2x concentration for 15 minutes at 37 °C. Stained cells were washed twice with PBS containing 2% FBS. Keratinocytes were identified as CD45^−^FAP^−^CD90^−^CD49f^+^CD29^+^ cells, melanocytes as CD45^−^FAP^−^CD90^−^CD49f^−^CD29^−^CD117^+^ cells, endothelial cells as CD45^−^EpCAM^−^CD31^+^ cells, FAP^+^ fibroblasts as CD45^−^CD31^−^FAP^+^ cells, and FAP^−^ fibroblasts as CD45^−^CD31^−^FAP^−^CD90^+^ cells[23]. For analysis, cells were fixed and analyzed by flow cytometry (BD FACSymphony A5). For cell sorting, stained cells were sorted using a BD FACSAria Fusion (BD Biosciences) equipped with a 100 µm nozzle.

### Bulk RNA-seq

For bulk RNA-seq, up to 400 cells were sorted directly into lysis buffer, and cDNA was generated using the SMART-Seq v4 Ultra Low Input RNA Kit for Sequencing (Takara). Libraries were prepared using the Nextera XT DNA sample preparation kit (Illumina). Samples were sequenced on an Illumina NextSeq 2000 using paired-end reads with a read length of 59 nt, and a sequencing depth of approximately 5 million reads per sample. Base-calling and demultiplexing were performed automatically on BaseSpace (Illumina) to generate FASTQ files. The FASTQs were aligned to the University of California Santa Cruz (UCSC) Human genome assembly version 19, using STAR v.2.4.2a and gene counts were generated using htseq-count. QC and metrics analysis was performed using the Picard family of tools (v1.134).

### Bulk ATAC-seq

ATAC-seq was performed following previously described protocols [24–26].Epidermal cell suspension was treated with 200 U/mL of DNaseI (Worthington Biochemical Corporation) at 37 °C for 10 minutes. Keratinocytes were then enriched from epidermal cell suspensions by depletion of CD45^+^ and CD117^+^ cells and removal of dead cells using magnetic bead-based separation. Cells were labeled with anti-PE MicroBeads (Miltenyi Biotec) following antibody staining and with Dead Cell Removal MicroBeads (Miltenyi Biotec), and subjected to a single magnetic column separation step. Flow through enriched for keratinocytes was collected. A total of 50,000 cells were used for nuclei isolation and tagmentation. Nuclei preparation and transposition reactions were performed following the Omni-ATAC-seq protocol as previously described [24, 25]. Briefly, cells were lysed in a lysis buffer containing IGEPAL CA-630, Tween 20, and digitonin, and nuclei were incubated with Tn5 transposase at 37 °C for 30 minutes. Transposed DNA was purified and amplified by PCR following Buenrostro et al [26]. Libraries were sequenced on an Illumina NextSeq 2000 using paired-end reads with a read length of 59 nt and a sequencing depth of 50-71.7 million reads per sample.

### RNA-seq data analysis

RNA-seq libraries were quality-controlled based on read counts, alignment rate, and gene coverage, and samples were excluded based on predefined thresholds (minimum 0.5 million reads, >75% alignment, and median coverage CV < 0.75). Gene-level counts were restricted to protein-coding genes with HGNC symbols, and lowly expressed genes were filtered based on a threshold of at least one count per million reads in at least 2% of samples. Counts were normalized using the TMM method, and differential expression analysis was performed using the limma-voom pipeline with donor included as a blocking factor. Analyses were performed separately for each cell type, and genes with FDR < 0.05 and |log2 fold change| > 1 were considered significant. Reactome pathway over-representation analysis (ORA) was performed separately for upregulated and downregulated genes using the ReactomePA package. All genes tested in the corresponding differential expression analysis were used as the background gene universe. P values were adjusted for multiple testing using the Benjamini-Hochberg method, and pathways with FDR < 0.05 were considered significantly enriched. For analysis of common IFN-γ-responsive pathways, genes upregulated in all five cell types at each culture time point were combined across time points to generate a unique common upregulated gene set for Reactome ORA. Similarly, cell-type-specific upregulated and downregulated genes were combined across time points for each cell type to generate unique gene sets for Reactome ORA. Hallmark IFN-γ response genes were obtained from the Molecular Signatures Database (MSigDB) Hallmark gene set collection using the msigdbr package. For each cell type and culture time point, the mean log2 fold change between IFN-γ-stimulated and unstimulated samples was calculated across Hallmark IFN-γ response genes without filtering for statistical significance. For ssGSEA, sample-level enrichment scores for the Hallmark IFN-γ-response gene set were calculated from log2-transformed normalized CPM values using the GSVA package. For comparison of cytokine responses between 2D-cultured and biopsy-derived keratinocytes, analyses were restricted to genes represented in both datasets.

### ATAC-seq data analysis

Peaks were called using MACS3 with ATAC-seq parameters. Quality control metrics, including fragment size distribution, transcription start site (TSS) enrichment, read depth, alignment rate, and fraction of reads in peaks (FRiP), were assessed based on ENCODE guidelines (https://www.encodeproject.org/atac-seq/). Peaks were called per library with MACS2 (–f BAMPE –g hs –q 0.001), then concatenated, filtered against the ENCODE hg38 blacklist, and merged into a consensus set. Peak-level accessibility was quantified as bigWig signal across the consensus peak set. Peak signals were normalized using a median of ratios normalization approach. Differential chromatin accessibility was assessed using linear modeling implemented in limma with voomWithQualityWeights. Regions with FDR < 0.05 and |log2 fold change| > 1 were considered differentially accessible. Peaks were annotated to the nearest gene and to promoters within ±10 kb using ChIPpeakAnno. Motif enrichment analysis was performed using HOMER [27] on differentially accessible regions with a specified background set and region size of 200 bp.

## Results

### Viable skin structural cells can be isolated from previously frozen biopsies for analysis

To establish an *ex vivo* culture system to study responses of human skin structural cells based on frozen skin biopsies, we first examined whether viable cells could be isolated from these samples. Biopsies were either processed immediately after thawing or cultured at an air liquid interface with the dermal side placed on 3.0 µm pore size Transwell inserts. We harvested biopsies for analysis by flow cytometry and RNA-seq at day 0 or after 1, 3, 5, 7, 10 or 14 days of culture, and cells were separately isolated from either the epidermis or dermis (Supplementary Fig. 1a). Among epidermal cells, keratinocytes were defined as CD45^−^ FAP^−^CD90^−^CD49f^+^CD29^+^ cells, and melanocytes as CD45^−^FAP^−^CD90^−^CD49f^−^CD29^−^CD117^+^ cells, whereas in the dermis endothelial cells were defined as CD45^−^EpCAM^−^CD31^+^ cells, FAP^+^ papillary fibroblasts as CD45^−^EpCAM^−^CD31^−^FAP^+^ cells, and FAP^−^ reticular fibroblasts as CD45^−^EpCAM^−^CD31^−^FAP^−^CD90^+^ cell (Fig. 1a, b).

**Figure 1.**
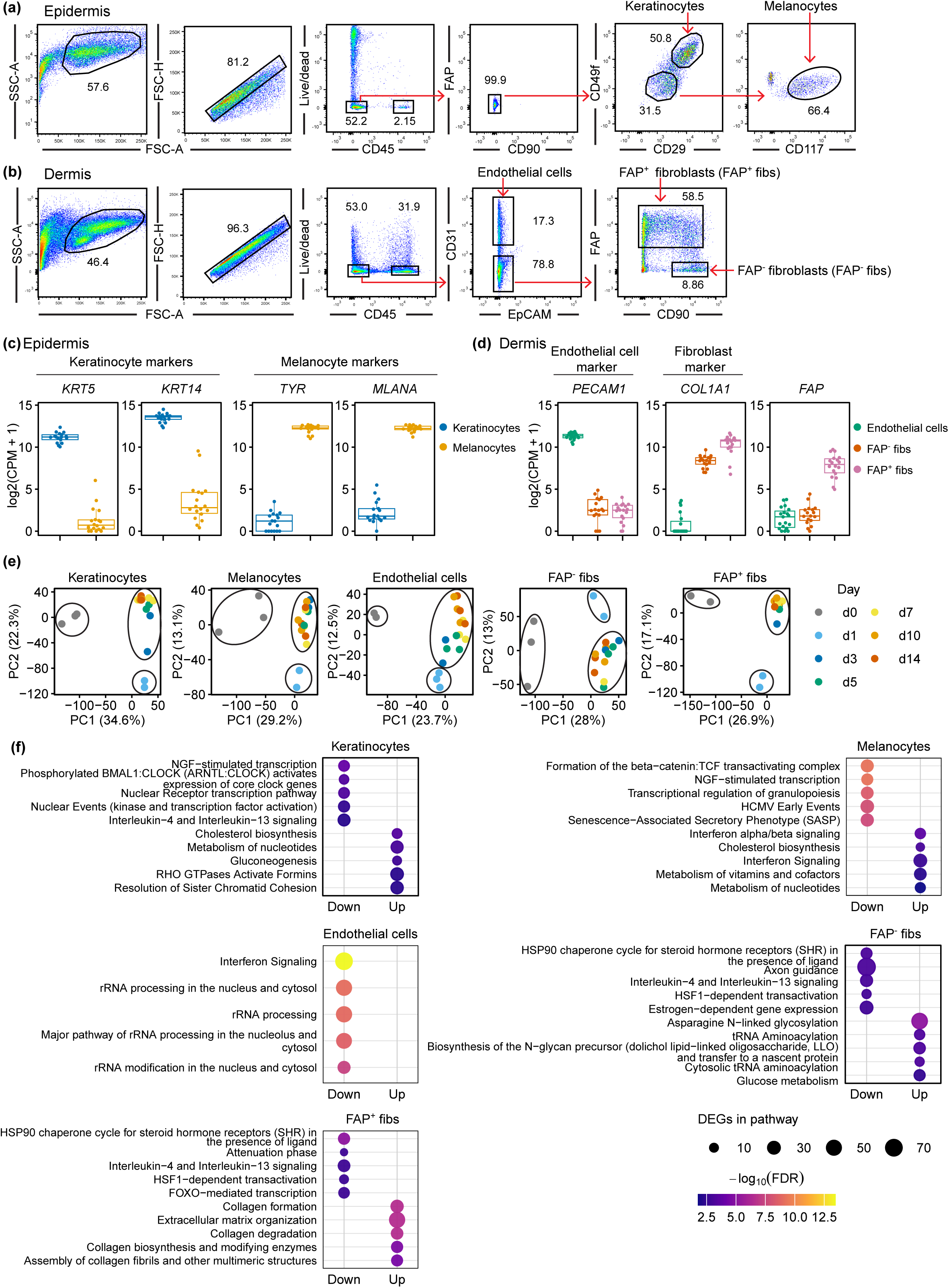
Skin structural cells isolated from frozen skin biopsies retain cell-type-specific transcriptional identities and exhibit distinct transcriptional responses during culture. Frozen skin biopsies from healthy donors were thawed and either processed immediately (day 0) or cultured for 1, 3, 5, 7, 10, or 14 days. At each time point, skin biopsies were processed to separate the epidermis and dermis, followed by sorting of keratinocytes melanocytes, endothelial cells, FAP-negative fibroblast (FAP^−^ fibs), and FAP-positive fibroblasts (FAP^+^ fibs) for bulk RNA-sequencing (RNA-seq). (a, b) Gating strategy for isolating skin structural cells from skin biopsies. Keratinocytes and melanocytes were isolated from the epidermis (a), whereas endothelial cells, FAP^−^ fibroblasts, and FAP^+^ fibroblasts were isolated from the dermis (b). (c, d) Normalized expression of the indicated cell type-specific genes in keratinocytes and melanocytes (c), and in endothelial cells, FAP^−^ fibroblasts, and FAP^+^ fibroblasts (d). (e) Principal component analysis (PCA) of bulk RNA-seq data performed separately for each cell type at day 0 and the indicated culture time points. (f) Reactome pathway enrichment analysis of genes differentially expressed between day 10 and day 0 in each cell type. Upregulated and downregulated genes (|log2 fold change| > 1, FDR < 0.05) were analyzed separately. Up to five significant Reactome pathways (FDR < 0.05) with the lowest FDR are shown for each direction and cell type. Dot size indicates the number of differentially expressed genes in each pathway, and color indicates –log10(FDR). RNA-seq analyses in (c-f) used unstimulated samples from three donors. The exact number of donors with QC-passing data for each cell type and day is provided in Supplementary Table 1.

To verify the identity of the sorted populations, we performed principal component analysis (PCA) and analyzed the expression of known lineage-specific marker genes. PCA showed that keratinocytes and melanocytes formed distinct clusters in the epidermis (Supplementary Fig. 1b), and in the dermis we observed three distinct clusters consistent with endothelial cells, FAP^+^ fibroblasts, and FAP^−^ fibroblasts (Supplementary Fig. 1c). Confirming the cell identities, keratinocyte marker genes such as *KRT5* and *KRT14* were preferentially expressed in CD45^−^FAP^−^CD90^−^CD49f^+^CD29^+^ cells, whereas melanocyte marker genes such as *TYR* and *MLANA* were specifically enriched in CD45^−^FAP^−^CD90^−^CD49f^−^CD29^−^CD117^+^ cells (Fig. 1c). Similarly, in the dermis the endothelial marker *PECAM1* was highly expressed in sorted CD45^−^ EpCAM^−^CD31^+^ cells, the fibroblast marker *COL1A1* was enriched in both CD45^−^EpCAM^−^CD31^−^FAP^+^ and as CD45^−^EpCAM^−^CD31^−^FAP^−^CD90^+^ cell populations, and *FAP* was enriched in FAP^+^ CD45^−^EpCAM^−^CD31 cells (Fig. 1d). These data demonstrate that the sorted populations were correctly identified and retained cell-type-specific transcriptional identities even during prolonged explant culture (Fig. 1c, d).

We next examined transcriptional changes that occurred during culture. PCA showed that in all cell types examined, samples processed immediately after thawing (day 0) formed a distinct cluster from cultured samples, and that gene expression largely stabilized after approximately 3 days of culture (Fig. 1e). Similarly, the number of differentially expressed genes (DEGs) changed markedly between day 0 and day 1 with subsequent changes becoming smaller and largely stabilizing by day 3 in most cell types and by day 5 in melanocytes (Supplementary Fig. 1d). Reactome pathway enrichment analysis of DEGs between day 0 and day 1 revealed that mitochondrial translation-related pathways were ranked among the top enriched pathways for upregulated genes in melanocytes, endothelial cells, FAP^−^ fibroblasts, and FAP^+^ fibroblasts during the early phase of culture (Supplementary Fig. 1e).

To further characterize transcriptional changes that occur during culture, we compared gene expression between day 0 and day 10 when gene expression profiles had largely stabilized. In keratinocytes, differentiation-related pathways were enriched among downregulated genes, whereas cell proliferation-related pathways were enriched among upregulated genes (Fig. 1f). In melanocytes, pathways such as Formation of the beta-catenin:TCF transactivating complex and NGF-stimulated transcription were enriched among downregulated genes, whereas interferon signaling and metabolism-related pathways were enriched among upregulated genes (Fig. 1f). In endothelial cells, interferon signaling and rRNA processing-related pathways were enriched among downregulated genes, but no pathways were significantly enriched among upregulated genes (Fig. 1f). Finally, FAP^−^ fibroblasts and FAP^+^ fibroblasts shared several pathways enriched among downregulated genes, including HSP90 chaperone cycle, Interleukin-4 and Interleukin-13 signaling, and HSF1-dependent transactivation, while distinct pathways were enriched among upregulated genes including tRNA aminoacylation– and N-linked glycosylation-related pathways in FAP^−^ fibroblasts and collagen-related pathways enriched in FAP^+^ fibroblasts (Fig. 1f). Taken together these results indicate that viable skin structural cells can be recovered from previously frozen cultured skin biopsies, and that different cell types exhibit distinct transcriptional responses during culture that stabilize after approximately 3 days.

### Skin structural cells in previously frozen biopsies respond to inflammatory cytokines

We next examined whether skin structural cells isolated from frozen skin biopsies can respond to inflammatory cytokine stimulation during *ex vivo* culture. For this, skin biopsies were treated with IFN-γ for 24 hours prior to harvest at each time point, followed by isolation of skin structural cells and transcriptional analysis by bulk RNA-seq (Fig. 2a). IFN-γ-responsive genes, including *CD74*, *HLA-DRA*, *CXCL9*, *CXCL10*, and *CXCL11*, were strongly upregulated following IFN-γ stimulation in skin structural cells (Fig. 2b). Indeed, IFN-γ-induced transcriptional changes were observed in all cell types throughout the culture period, although the number of DEGs tended to be lower at day 1 than at later time points in most cell types (Fig. 2c). Consistent with this observation, the mean log2 fold change of Hallmark IFN-γ response genes tended to increase from day 1 to day 3 and remained relatively higher and became more consistent after day 5 of culture (Fig. 2d). To further characterize the differences in IFN-γ responses between day 1 and day 5, we compared the sets of DEGs identified at these two time points. The sets of significantly IFN-γ-responsive genes differed between day 1 and day 5 in some cell types. Among genes that were significantly responsive at both time points, the magnitude of IFN-γ-induced gene expression tended to be greater at day 5 than at day 1 (Supplementary Fig.2a)

**Figure 2.**
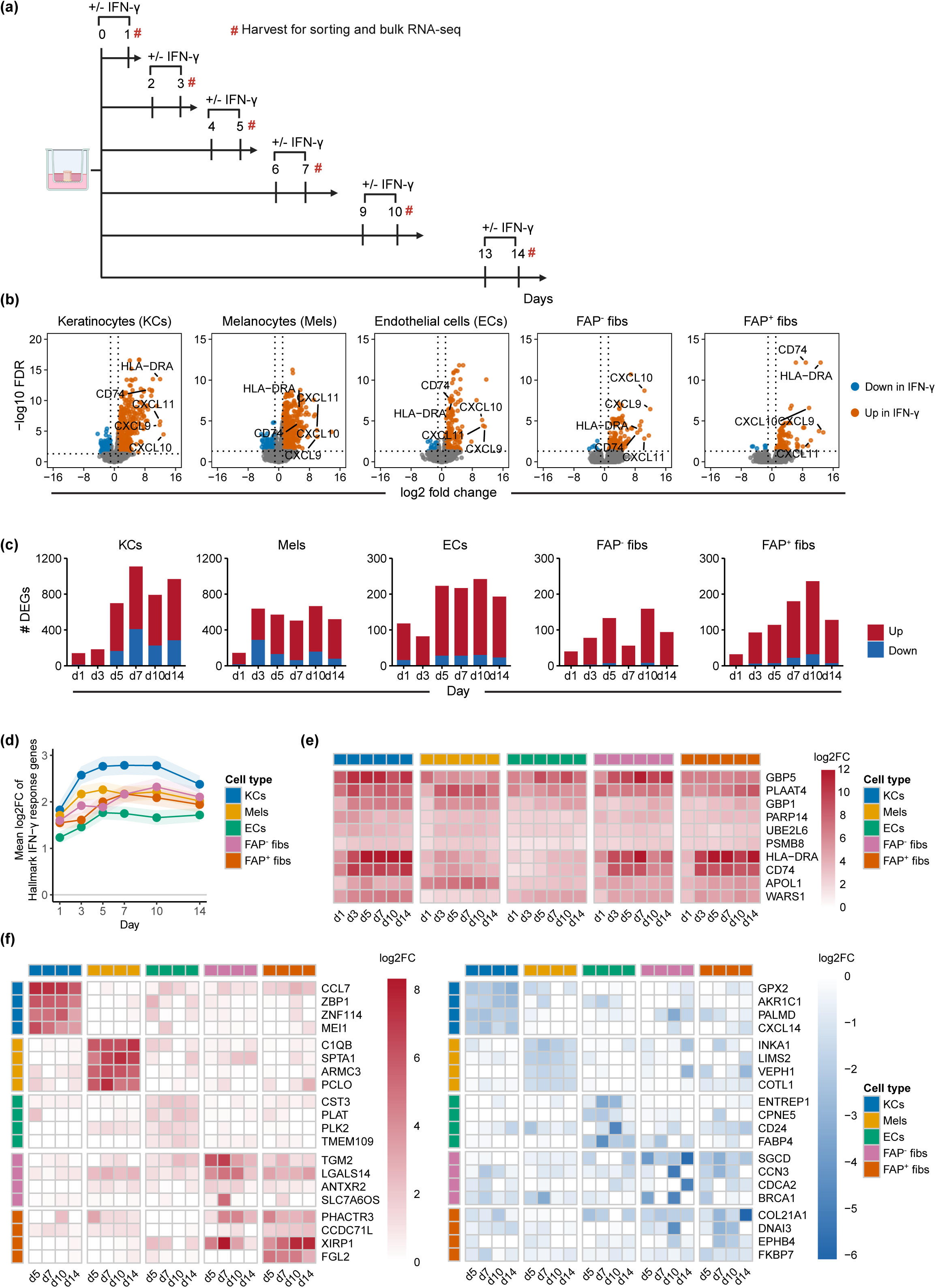
Skin structural cells can respond to IFN-γ during *ex vivo* culture. Frozen skin biopsies from healthy donors were cultured *ex vivo* with or without IFN-γ at indicated culture time points. Skin structural cell populations were subsequently isolated by cell sorting and subjected to bulk RNA-seq to assess their transcriptional response to IFN-γ. (a) Experimental design for investigating IFN-γ responses in skin structural cells. (b) Volcano plots of differential gene expression between IFN-γ-stimulated and unstimulated samples at day 5 in keratinocytes (KCs), melanocytes (Mels), endothelial cells (ECs), FAP^−^ fibroblasts, and FAP^+^ fibroblasts. Blue and orange dots indicate genes significantly downregulated and upregulated by IFN-γ, respectively, whereas gray dots indicate non-significant genes. Dotted lines indicated the threshold for differential expression (|log2 fold change| >1, FDR < 0.05). Selected genes are labeled. (c) Number of differentially expressed genes (DEGs) between IFN-γ-stimulated and unstimulated samples at each culture time points in the indicated cell types. DEGs were defined as genes with |log2 fold change| > 1 and FDR < 0.05. Blue and red bars indicate down regulated and upregulated genes, respectively. (d) Mean log2 fold change of Hallmark IFN-γ response genes across culture time points in the indicated cell types. Mean log2 fold change was calculated between IFN-γ-stimulated and unstimulated samples at each time point without filtering for statistical significance. Shaded area indicate the standard error of the mean and the gray horizontal line indicates no change between IFN-γ-stimulated and unstimulated samples. (e) Heatmap showing the top 10 genes commonly differentially expressed in response to IFN-γ across the five cell types. Common DEGs were defined based on differential expression between IFN-γ-stimulated and unstimulated samples at each culture time point (|log2 fold change| > 1, FDR < 0.05). Genes were ranked by the number of culture time points at which they were identified as common DEGs, followed by mean log2 fold change. Heatmap values indicate log2 fold change between IFN-γ-stimulated and unstimulated samples. (f) Heatmap showing the top four cell-type-specific genes differentially expressed between IFN-γ-stimulated and unstimulated samples in each cell type from day 5 to day 14 (left, upregulated; right, downregulated). Cell-type-specific DEGs were defined as genes differentially expressed in one cell type (|log2 fold change| > 1, FDR < 0,05) but not differentially expressed in other cell types at the same culture time point. Genes were ranked by the number of culture time points at which they were identified as cell type-specific DEGs, followed by mean log2 fold change with upregulated and downregulated genes ranked separately. Heatmap values indicate log2 fold change between IFN-γ-stimulated and unstimulated samples. RNA-seq analyses in (b-f) used unstimulated and IFN-γ-stimulated samples from three donors. The number of donors with QC-passing data for each cell type, condition, and culture day is provided in Supplementary Table 1.

We next examined common and cell-type-specific IFN-γ-responsive DEGs among the five cell types during *ex vivo* culture. The number of DEGs commonly upregulated by IFN-γ across the five cell types ranged from 10 to 78 across the culture time points, whereas no DEGs were commonly downregulated across the five cell types (Supplementary Fig. 2b). Among the top 10 common DEGs were IFN-γ responsive genes including guanylate-binding proteins (GBPs), *HLA-DRA*, and *CD74* (Fig. 2e). These common upregulated genes were enriched in interferon signaling and antigen processing-related pathways (Supplementary Fig. 2d). The number of both upregulated and downregulated cell-type-specific DEGs was higher in keratinocytes and melanocytes than in endothelial cells, FAP^−^ fibroblasts, and FAP^+^ fibroblasts (Supplementary Fig. 2d). Distinct sets of genes were upregulated and downregulated by IFN-γ in a cell-type-specific manner (Fig. 2f) Reactome pathway analysis further revealed distinct patterns among the cell types. Keratinocyte-specific upregulated genes were enriched in interferon-related signaling pathways, whereas downregulated genes were enriched in cell-cycle-related pathways. Melanocyte-specific upregulated genes were predominantly enriched in TLR-related signaling pathways, whereas downregulated genes were enriched in pathways to keratinization and keratinocyte-differentiation. Endothelial cell-specific upregulated genes were enriched in NLR-related and immune signaling pathways, while FAP^+^ fibroblast-specific downregulated genes were enriched in MET-and extracellular matrix-related pathways. No significant pathway enrichment was observed for remaining cell type-specific gene sets (Supplementary Fig. 2e). Taken together, these data indicate that skin structural cells isolated from frozen skin biopsies retain cytokine responsiveness and that IFN-γ-responsive gene expression becomes more consistent after day 5 of culture. IFN-γ-induced transcriptional responses were both common across cell types and cell-type specific.

We further examined whether skin structural cells derived from frozen skin biopsies can respond to additional inflammatory cytokines. Biopsies were cultured for 5 days and then stimulated with IFN-γ, TNF-α, IL-17A, IL-13, or IL-1β for 24 hours prior to harvest. After digestion, keratinocytes and melanocytes were sorted for bulk RNA-seq (Supplementary Fig. 3a). In keratinocytes TNF-α induced *ICAM1* and *CCL20*, IL-17A induced *DEFB4A*, *S100A7*, and *LCN2*, IL-13 induced *CCL26, SERPINB3* and *SERPINB4*, and IL-1β induced *LCN2* and *CCL20*. In melanocytes TNF-α induced *CXCL8* and *TNFAIP6*, whereas IL-1β induced *FDCSP, CCL5*. IL-13 also induced transcriptional changes, although the genes showing the largest increases did not include canonical IL-13 responsive genes. Furthermore IL-17A elicited a much more limited transcriptional response (Supplementary Fig. 3b). Together these results indicate that skin structural cells retain the ability to respond to multiple cytokine stimuli in cultured freeze-thawed skin biopsies with distinct transcriptional responses across cell types. We further compared cytokine responses in keratinocytes with those of 2D-cultured immortalized keratinocytes to examine whether the response patterns differed between the two culture conditions. For IFN-γ stimulation, the magnitude of gene expression changes tended to be greater in keratinocytes isolated from skin biopsies, while the responses were positively correlated between the two conditions (Pearson *r* = 0.55, Spearman *ρ* = 0.40). In contrast, IL-17A and IL-13 induced gene expression changes tended to be greater in 2D-cultured keratinocytes, and correlations of the response between the two conditions were weak for both IL-17A (Pearson *r* = 0.16, Spearman *ρ* = 0.08) and IL-13 (Pearson *r* = 0.16, Spearman *ρ* = 0.15) (Supplementary Fig. 3c). These observations suggest that the tissue context may influence cytokine-induced transcriptional responses in keratinocytes.

### Rapid resolution of IFN-γ responses in human skin structural cells

To determine whether explant cultures can be used to study resolution of inflammatory responses, we next examined transcriptional responses after stimulus removal. For IFN-γ-removal condition, biopsies were stimulated with IFN-γ on day 4 for 24 hours, washed on day 5 and cultured for additional 5 days in the absence of IFN-γ. On day 10, biopsies were processed, and keratinocytes, melanocytes, and endothelial cells were sorted for bulk RNA-seq (Fig. 3a). Comparing IFN-γ-stimulated/rested samples with unstimulated samples cultured for 10 days, no significantly differentially expressed genes were detected in either keratinocytes or endothelial cells. In melanocytes, however, a small number of genes remained differentially expressed and these were IFN-γ-responsive genes, indicating incomplete resolution of the IFN-γ responses (Fig. 3b). Consistent with these findings, Hallmark IFN-γ response scores in IFN-γ-removal samples at day 10 were comparable to those in unstimulated samples at day 10, whereas the scores were higher in IFN-γ-stimulated than in unstimulated samples at day 5 in all cell types examined (Fig. 3c). These data demonstrate that IFN-γ-induced transcriptional changes in these populations of skin structural cells were highly dynamic and largely resolved after IFN-γ removal.

**Figure 3.**
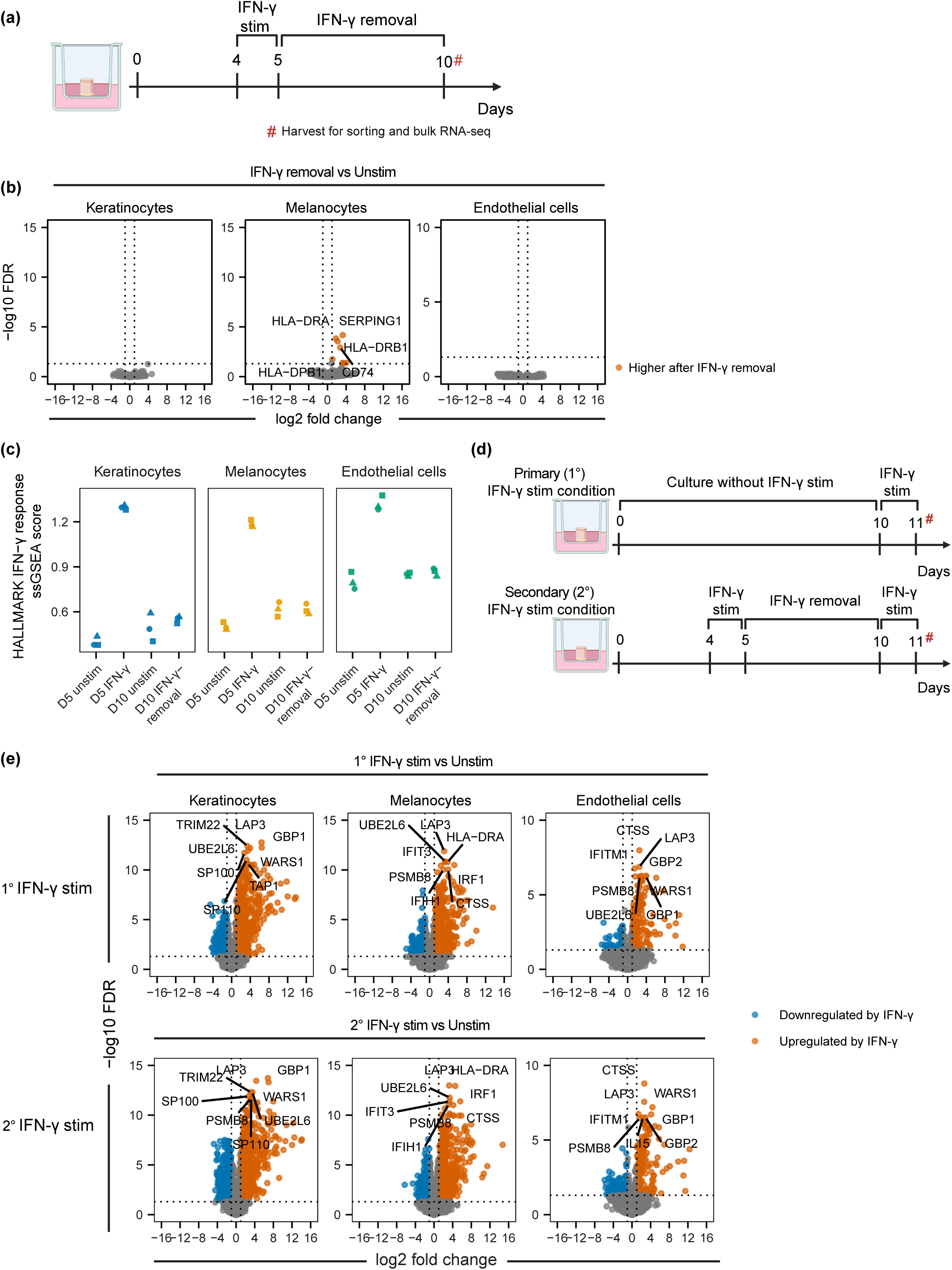
IFN-γ-induced transcriptional changes resolve after stimulus removal, and skin structural cells remain responsive to secondary IFN-γ stimulation. Frozen skin biopsies from healthy donors were cultured *ex vivo* and stimulated with IFN-γ, followed by stimulus removal to assess the resolution of IFN-γ-induced transcriptional changes. In a separate experiment, skin biopsies were subjected to primary (1°) or secondary (2°) IFN-γ stimulation to assess the transcriptional response to secondary IFN-γ stimulation. Skin structural cell populations were isolated by cell sorting and subjected to bulk RNA-seq. (a) Experimental design for assessing the resolution of IFN-γ-induced transcriptional changes after stimulus removal. (b) Volcano plots of gene expression in keratinocytes, melanocytes, and endothelial cells comparing IFN-γ removal samples with unstimulated samples cultured for 10 days. Orange dots indicate genes with significantly higher expression following IFN-γ removal, whereas gray dots represent non-significant genes. Dotted lines indicate the thresholds for differential expression (|log2 fold change| > 1, FDR < 0.05). (c) Hallmark IFN-γ response ssGSEA scores in keratinocytes, melanocytes, and endothelial cells from unstimulated samples at day 5 (D5 unstim), IFN-γ-stimulated samples at day 5 (D5 IFN-γ), unstimulated samples at day 10 (D10 unstim), and d10 samples after IFN-γ removal (D10 IFN-γ removal). Each symbol represents an individual donor (d) Experimental design for primary (1°) and secondary (2°) IFN-γ stimulation in skin structural cells. (e) Volcano plots of differential gene expression in keratinocytes, melanocytes, and endothelial cells comparing primary (1°) IFN-γ-stimulated samples (top) or secondary (2°) IFN-γ-stimulated samples (bottom) with unstimulated samples cultured for 10 days. Blue and orange dots indicate genes significantly downregulated and upregulated respectively, compared with unstimulated samples, whereas gray dots indicate not-significant genes. Dotted lines indicate the thresholds for differential expression (|log2 fold change| > 1, FDR < 0.05). RNA-seq was performed using samples n = 3 independent donors per cell type and condition.

We next examined whether skin structural cells respond to secondary IFN-γ stimulation and whether there were any changes in a secondary response indicative of inflammatory memory. Following IFN-γ stimulation and washout, biopsies were cultured without stimulation for an additional 5 days, subjected to secondary IFN-γ stimulation for 24 hours on day 10 and processed for bulk RNA-seq (Fig. 3d). Secondary stimulation induced robustly upregulated IFN-γ-responsive genes in all cell types compared with unstimulated samples (Fig. 3e). Primary IFN-γ stimulation at day 10 also induced IFN-γ-responsive genes compared with unstimulated samples (Fig. 3d, e), and IFN-γ-induced gene expression changes were highly correlated between primary and secondary IFN-γ stimulation across these cell types (Supplementary Fig. 4a). Direct comparison of primary and secondary responses identified differences in a limited number of genes, including *L3MBTL3* and *HOPX* in keratinocytes, *ID2* and *P2RX7* in melanocytes, and *STON2* and *OR52H1* in endothelial cells. (Supplementary Fig. 4b). Collectively, these data indicate that IFN-γ-induced transcriptional changes largely resolve after stimulus removal, and that although skin structural cells remain responsive to IFN-γ secondary stimulation, they do not exhibit altered responses indicative of inflammatory memory upon prior IFN-γ stimulation.

### Chromatin remodeling and resolution following IFN-γ stimulation in keratinocytes

We next investigated whether the skin explant culture system could be used to monitor changes in chromatin accessibility in skin structural cells during inflammatory cytokine treatment using bulk ATAC-seq from isolated keratinocytes. For this, biopsies were stimulated with IFN-γ on day 4 for 24 hours, and keratinocytes were enriched for bulk ATAC-seq using bead-based enrichment to reduce nuclei damage associated with cell sorting (Fig. 4a, Supplementary Fig. 5a). Sample quality was assessed by fragment size distribution, TSS enrichment, FRiP score, and library size. Fragment size distribution showed enrichment of nucleosome-free fragments (< 100 bp) and clear mono-nucleosome peaks. TSS enrichment profiles showed the expected enrichment around transcription start sites, FRiP scores ranged from 0.47 to 0.65, and estimated library complexity ranged from 54.4 to 102.9 million unique fragments (Supplementary Fig. 5b), indicating high quality of tagmentation and library construction. Consistent with the robust transcriptional responses we observed, IFN-γ treatment substantially altered the chromatin accessibility landscape of keratinocytes, with both significant increases and decreases in chromatin accessibility at different loci, including known IFN-γ responsive genes such as *CXCL10* and *CXCL11* (Fig. 4b, c). Motif analysis further revealed enrichment of interferon-stimulated response element (ISRE) and IRF family motifs in IFN-γ-induced open chromatin regions (Fig. 4d), in line with IFN-γ-induced transcriptional changes observed in RNA-seq. In contrast, regions showing decreased chromatin accessibility were enriched for *GRHL2, GATA, and NRF2* motifs (Fig. 4d). In keratinocytes, GRHL2 and GATA3 both modulate terminal differentiation [28], [29], whereas NRF2 regulates antioxidant defense [30].

**Figure 4.**
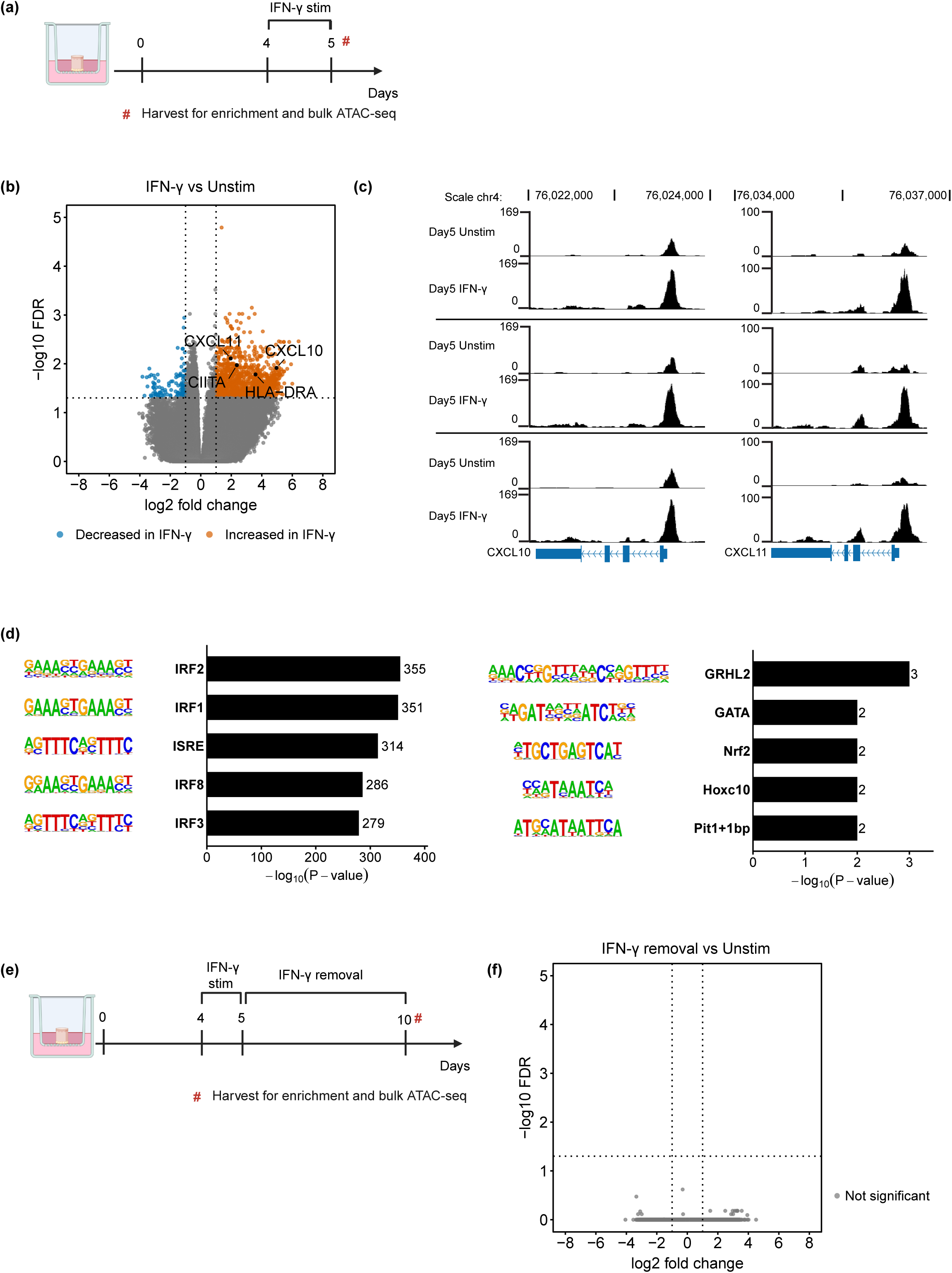
IFN-γ-induced changes in chromatin accessibility are associated with IFN-γ-responsive regulatory elements and resolve after stimulus removal. Frozen skin biopsies from healthy donors were cultured *ex vivo* with or without IFN-γ stimulation. Keratinocytes were subsequently enriched and subjected to bulk ATAC-seq to assess IFN-γ-induced changes in chromatin accessibility. (a) Experimental design for investigating IFN-γ-induced changes in chromatin accessibility in keratinocytes. (b) Volcano plots of IFN-γ-induced changes in chromatin accessibility at day 5. Blue and orange dots indicate regions with significantly decreased and increased accessibility, respectively, in IFN-γ-stimulated samples compared with unstimulated samples in keratinocytes. Dotted lines indicate the thresholds for differential accessibility (|log2 fold change| > 1, FDR < 0.05). (c) UCSC Genome Browser tracks showing chromatin accessibility at *CXCL10* (left) and *CXCL11* (right) loci in keratinocytes at day 5. Tracks show paired unstimulated and IFN-γ-stimulated samples from the same donors. (d) Motif enrichment analysis was performed using HOMER. Differentially accessible regions with increased and decreased accessibility following IFN-γ stimulation were analyzed using a specific background set and a region size of 200 bp. (e) Experimental design for examining IFN-γ-induced changes in chromatin accessibility after IFN-γ removal. (f) Volcano plots comparing changes in chromatin accessibility in keratinocytes between samples following IFN-γ removal and samples cultured for 10 days without stimulation. ATAC-seq was performed using samples from n = 3 independent donors per condition.

To further determine how changes in chromatin accessibility resolved after IFN-γ stimulus removal, biopsies from 3 individual donors were stimulated with IFN-γ for 24 hours on day 4, washed on day 5 and cultured for an additional 5 days without stimulation. On day 10, biopsies were harvested and keratinocytes were isolated for bulk ATAC-seq (Fig. 4e). When comparing IFN-γ-removal samples with unstimulated samples cultured for 10 days, IFN-γ-induced chromatin accessibility changes were almost completely resolved after stimulus removal (Fig. 4f). These results are consistent with the rapid resolution of IFN-γ-induced transcriptional changes in keratinocytes observed by RNA-seq, and the lack of inflammatory memory in transcriptional responses upon secondary stimulation.

### Inhibition of class 1 HDACs or EZH2 do not impact transcriptional resolution of IFN-γ responses

Next, we mechanistically explored the rapid resolution of transcriptional and epigenetic changes induced by IFN-γ stimulation. We focused on histone deacetylases (HDACs), which remove acetyl groups from histones [31] and EZH2, which catalyzes trimethylation of histone H3 at lysine 27 [32]. Entinostat, an inhibitor of multiple class I HDACs, has been reported to increase the expression of a subset of IFN-γ-responsive genes in macrophages [33]. Tazemetostat, EZH2 inhibitor has shown to restore IFN-γ-induced expression of MHC-class II-related-genes in melanoma cell lines [34]. We therefore used these inhibitors to examine whether HDACs and EZH2 contribute to resolution of IFN-γ transcriptional responses. Biopsies were cultured and treated with IFN-γ for 24 hours on day 4. On day 5, immediately after IFN-γ removal, the biopsies were treated with entinostat, tazemetostat or DMSO as vehicle control. The treatment medium was replaced every other day through day 9. On day 10, biopsies were harvested and keratinocytes were sorted for bulk RNA-seq (Fig. 5a). Flow cytometry analysis confirmed that entinostat increased H3K27ac levels, while tazemetostat decreased H3K27me3 levels in keratinocytes from inhibitor-treated samples compared with DMSO-treated controls (Fig. 5b). Entinostat treatment induced significantly upregulated genes compared with DMSO-treated controls under both unstimulated and IFN-γ-removal conditions. However, only one gene, *POTEE*, was detected as differentially expressed genes when comparing the effect of entinostat relative to DMSO between IFN-γ-removal and unstimulated samples. *POTEE* belongs to a highly paralogous family in which short reads are difficult to assign to the correct paralog, and we therefore considered this a likely technical artifact. These results suggest that entinostat induce transcriptional changes in keratinocytes, but that these changes do not impact the resolution of IFN-γ responses (Fig. 5c). Furthermore, tazemetostat treatment did not induce significantly differentially expressed genes across any comparisons, even though tazemetostat reduced H3K27me3 levels (Fig. 5c). Hallmark IFN-γ response scores were similar between unstimulated samples and IFN-γ-removal samples on day 10 across DMSO– and inhibitor-treated group, whereas scores were higher in IFN-γ-stimulated samples than in unstimulated samples on day 5 (Fig. 5d). These data suggest that neither class 1 HDACs nor EZH2 were central to the transcriptional resolution of IFN-γ responses.

**Figure 5.**
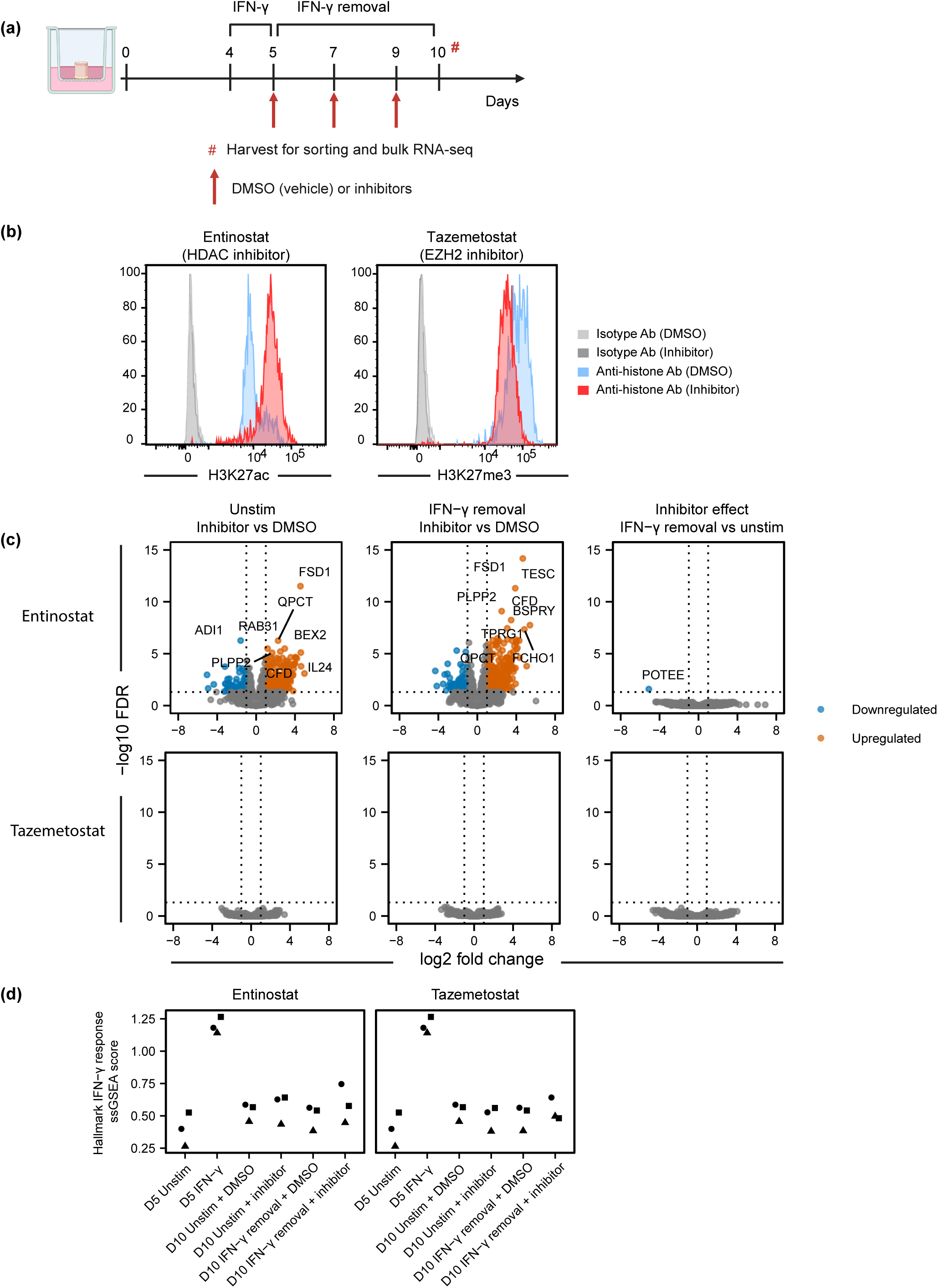
Histone-modifying enzyme inhibitors do not substantially affect transcriptional resolution IFN-γ responses in keratinocytes. Frozen skin biopsies from healthy donors were cultured *ex vivo* and stimulated with IFN-γ, followed by IFN-γ removal and treatment with histone-modifying enzyme inhibitors during the resolution period. Keratinocytes were subsequently isolated and subjected to bulk RNA-seq. (a) Experimental design for investigating the effects of histone-modifying enzyme inhibitors on transcriptional resolution after IFN-γ removal in keratinocytes. (b) Flow cytometric analysis of H3K27ac and H3K27me3 levels in keratinocytes isolated from frozen skin biopsies treated with entinostat or tazemetostat as indicated. (c) Volcano plots of differentially expressed genes in keratinocytes isolated at day 10 from frozen skin biopsies treated with entinostat (top) or tazemetostat (bottom). The left and middle panels compare inhibitor-treated samples with DMSO-treated samples under unstimulated (left) and IFN-γ removal (middle) conditions, respectively. The right panels compare the inhibitor effect relative to DMSO between IFN-γ removal and unstimulated conditions. Blue and orange dots indicate genes significantly downregulated and upregulated, respectively, whereas gray dots indicate non-significant genes in each indicated comparison. Dotted lines indicate the threshold for differential expression (|log2 fold change| > 1, FDR < 0.05). Up to eight significant genes with the lowest FDR are labeled. (d) Hallmark IFN-γ response ssGSEA scores in keratinocytes isolated from frozen skin biopsies treated with entinostat (left) or tazemetostat (right) under the indicated conditions. RNA-seq analyses in (c,d) used samples from n = 3 independent donors per condition. Each symbol represents an individual donor.

## Discussion

In this study, we established an *ex vivo* culture system to assess transcriptional and chromatin accessibility changes in skin structural cells in response to immune stimuli using frozen human skin biopsies. We show that viable skin structural cells can be isolated from both uncultured and cultured freeze-thawed skin biopsies, and they retain cell-type-specific transcriptional profiles. Using IFN-γ stimulation, we demonstrated that our system can evaluate dynamic transcriptional and chromatin accessibility changes in skin structural cells. These changes largely resolved after cytokine removal while cells remained transcriptionally responsive to secondary stimulation. We further demonstrated that this system can be used to assess the effects of drug candidates such as those targeting histone-modifying enzymes.

Current platforms to assess cytokine responses in skin structural cells are limited in physiological relevance because they lack or only partially retain skin structure, often do not include all relevant skin structural cell types, and often rely on immortalized cell lines. Our system can evaluate cytokine responses in skin structural cells *ex vivo*, maintaining skin structural cell identities and primary cell composition during culture. Explant culture of skin biopsies has been used to assess responses to wounding or topical therapeutics, but these systems only used fresh tissue samples and did not evaluate responses of the individual cellular components of the skin [35, 36]. Use of frozen skin biopsies allows us to create sample banks and increase experimental flexibility since we can choose experimental timing compared with fresh skin samples, thereby reducing batch effects and improving reproducibility. Furthermore, our system can simultaneously assess cytokine responses in skin structural cells from multiple donors, better capturing individual biological variability and enabling response profiling in samples from various inflammatory skin diseases. Overall, our system provides a physiologically relevant and experimentally flexible platform for evaluating cytokine responses across multiple cell types, including melanocytes and endothelial cells, which are not generally represented *in vitro* skin models.

Baseline gene expression profiles of skin structural cells changed during initial culture, but rapidly stabilized after 3-5 days. This stabilization may provide a more reliable baseline for evaluating cytokine-induced transcriptional responses. In addition to IFN-γ, we also detected transcriptional responses to other inflammatory cytokines, including TNF-α, IL-17A and IL-13, supporting the utility of this system for profiling cytokine responses in skin structural cells. When comparing cytokine responses in keratinocytes from 2D cultures and skin biopsies, we observed that some genes changed in the same direction under both conditions, whereas the responses were poorly correlated following IL-17A or IL-13 stimulation, indicating key differences in response profiles in genuine tissue cells vs cultured cell lines.

We also evaluated IFN-γ-induced chromatin accessibility changes in keratinocytes. Although chromatin accessibility changes have been reported in skin structural cells of patients with chronic inflammatory skin diseases [10], the upstream signals, such as cytokines, that drive these disease-associated changes remain poorly defined. Since *ex vivo* systems for assessing chromatin accessibility changes in response to immune stimuli remain limited, our system can be used to mechanistically dissect disease-associated chromatin remodeling in skin structural cells. Furthermore, it has been reported that histone modifications are involved in the pathogenesis of chronic skin diseases and histone modification inhibitors are potential therapeutic targets for atopic dermatitis and psoriasis [37], and our system may be useful for evaluating candidate therapeutic compounds in skin structural cells.

Beyond its utility as a platform, our study provides several biological insights into IFN-γ responses in skin structural cells. First, we found that IFN-γ-induced transcriptional and chromatin accessibility changes were largely reversed following stimulus removal. This contrasts with previous findings in human macrophages, where these changes persisted after IFN-γ washout but were reversed by treatment with a neutralizing IFN-γ antibody or JAK inhibitor [38]. Thus, IFN-γ washout alone may be sufficient to terminate signaling in skin structural cells, allowing restoration of their baseline transcriptional and chromatin accessibility states.

Second, we found little evidence of transcriptional memory following IFN-γ exposure, as secondary stimulation induced robust transcriptional responses of similar magnitude to those observed following primary stimulation. This contrasts with previous studies demonstrating inflammatory memory in other cell types. In mouse epidermal stem cells, inflammation-induced chromatin accessibility persists after stimulus removal, accompanied by continued binding of JUN, ATF3, and p63 at these loci [14]. Similarly, prior inflammation in mouse colonic stem cells promotes subsequent tumor growth and is associated with persistent increases in chromatin accessibility at AP-1 binding sites [39]. Transcriptional memory of IFN-γ signaling has also been reported in HeLa cells, where specific histone modifications were maintained despite the resolution of chromatin accessibility changes [40, 41]. Although we did not directly assess transcription factor occupancy or histone modifications following IFN-γ removal, differences in the establishment or maintenance of these epigenetic features may explain the absence of detectable inflammatory memory in skin structural cells. The duration of inflammatory stimulation may also be important, as previous studies demonstrating inflammatory memory in epidermal stem cells employed repeated stimulation over more than five consecutive days [13]. Prolonged inflammatory exposure may therefore be required to establish sustained transcription factor occupancy or histone modifications that support lasting changes in chromatin state and subsequent transcriptional responses. Taken together, these observations suggest that the regulation of secondary responses, including inflammatory memory and transcriptional memory, may depend on the nature and duration of the stimulus, cell type and tissue context. Future experiments will allow us to investigate the dynamics of skin structural cell responses to repeated exposure to inflammatory cytokines, as observed in chronic inflammatory skin disease, thereby helping elucidate the mechanisms underlying chronic inflammatory skin diseases and development of inflammatory memory.

Several limitations of the current study should be considered. Because skin punch biopsies were cultured for up to 14 days to assess IFN-γ-induced transcriptional responses, it remains unclear whether IFN-γ-induced responses are maintained during longer periods of culture. As a proof of concept, we stimulated biopsies with IFN-γ at a single concentration for 24 hours. IFN-γ is a cytokine produced by T cells and NK cells, and it robustly induces transcriptional responses in skin structural cells. Because we did not assess multiple cytokine stimulations, longer stimulation durations or a combination of both, it remains unclear whether those conditions would affect the resolution or alter secondary responses following primary stimulation. In addition, to investigate the role of histone modifications in IFN-γ-induced transcriptional resolution, we used histone modification inhibitors. Only two inhibitors, entinostat, an HDAC inhibitor, and tazemetostat, an EZH2 inhibitor, were tested. The involvement of other histone-modifying enzymes cannot be excluded in cytokine-induced transcriptional resolution.

In conclusion, we established an *ex vivo* platform to analyze cytokine-induced transcriptional and epigenetic responses in skin structural cells using frozen human skin biopsies. This system retains the cellular diversity and physiological architecture of native skin while offering the experimental flexibility and reproducibility afforded by biobanked frozen samples, enabling multi-donor, multi-cytokine, and multi-cell-type profiling that is difficult to achieve with existing *in vitro* or explant models. By capturing both the induction and resolution of cytokine-driven transcriptional and chromatin states, this platform provides a tractable framework for dissecting the mechanisms that drive chronic inflammatory skin diseases and for evaluating candidate therapeutics, including epigenetic modulators, in a physiologically relevant context. Future application of this system across a broader range of cytokines, stimulation regimens, and patient-derived samples should help clarify how skin structural cells encode and respond to recurrent inflammatory signals, and may ultimately inform the development of targeted therapies for atopic dermatitis, psoriasis, and other chronic inflammatory skin conditions.

## Supplementary Figure legends

**Supplementary Figure 1.** Skin structural cells isolated from frozen skin biopsies retain cell-type-specific transcriptional identities and exhibit distinct transcriptional responses during culture. Frozen skin biopsies from healthy donors were thawed and either processed immediately (day 0) or cultured for the indicated times. At each time point, skin structural cell populations were isolated by cell sorting and subjected to bulk RNA-seq. (a) Experimental design for assessing cellular and transcriptional changes in cultured frozen human skin biopsies. (b, c) Principal component analysis (PCA) of bulk RNA-seq data from epidermal cell populations, including keratinocytes and melanocytes (b), and dermal cell populations including endothelial cells, FAP^−^ fibroblasts, and FAP^+^ fibroblasts (c). (d) Number of differentially expressed genes (DEGs) in each cell type between consecutive culture time points. DEGs were defined as genes with |log2 fold change| > 1 and FDR < 0.05. Blue and red bars indicate downregulated and upregulated, respectively. Numbers above the bars indicated the total number of DEGs. (e) Reactome pathway enrichment analysis of genes differentially expressed between day 1 and day 0 in each cell type. Upregulated and downregulated genes (|log2 fold change| > 1, FDR < 0.05) were analyzed separately. Up to five significant Reactome pathways (FDR < 0.05) with the lowest FDR are shown for each direction and cell type. Dot size indicates the number of differentially expressed genes in each pathway and color indicates –log10(FDR). RNA-seq analyses in (b-e) used unstimulated samples from three donors. The exact number of donors with QC-passing data for each cell type and day is provided in Supplementary Table 1.

**Supplementary Figure 2.** Skin structural cells respond to IFN-γ during *ex vivo* culture. Frozen skin biopsies from healthy donors were cultured *ex vivo* and stimulated with IFN-γ to assess the transcriptional responses of skin structural cells. (a) Scatter plots of IFN-γ-responsive genes in the indicated cell types comparing the responses at day 1 and day 5. Blue, red, and gray dots indicate genes differentially expressed only at day 1, only at day 5 and at both time points, respectively. Differentially expressed genes (DEGs) were defined as genes with |log2 fold change| > 1 and FDR < 0.05. The top five genes ranked by the sum of the absolute log2 fold changes at day 1 and day 5 are labeled. (b) Number of common upregulated differentially expressed genes (DEGs) at each culture time point. Common upregulated DEGs were defined as genes significantly upregulated by IFN-γ-stimulation in all five cell types at the same culture time points (log2 fold change > 1, FDR < 0.05). (c) Reactome pathway enrichment analysis of common upregulated DEGs identified across the culture time course. The top eight enriched pathways (FDR < 0.05), ranked by FDR are shown. (d) Heatmap showing the numbers of cell-type-specific upregulated (left) and downregulated (right) differentially expressed genes (DEGs) between IFN-γ-stimulated and unstimulated samples at each time point. Cell-type-specific DEGs were defined as genes significantly upregulated (log2 fold change > 1, FDR < 0.05) or downregulated (log2 fold change < –1, FDR < 0.05) in one cell type but not differentially expressed in the other cell types at the same culture time point. (e) Reactome pathway enrichment analysis of cell-type-specific DEGs identified across the culture time point, shown separately for upregulated (left) and downregulated (right) genes. Up to five significantly enriched pathways (FDR < 0.05) with the lowest FDR are shown for each cell type and direction. Cell types with no significantly enriched pathways are not shown. Dot size indicates the number of differentially expressed genes in each pathway and color indicates –log10(FDR). RNA-seq analyses used in unstimulated and IFN-γ-stimulated samples from three donors. The number of donors with QC-passing data for each cell type, condition, and culture day is provided in Supplementary Table 1.

**Supplementary Figure 3.** Skin structural cells isolated from frozen skin biopsies show transcriptional responses to multiple cytokines. Frozen skin biopsies from a healthy donor (n = 1) were cultured *ex vivo* with or without IFN-γ, TNF-α, IL-17A, IL-13, or IL-1β. Keratinocytes and melanocytes were subsequently isolated for assessment of transcriptional responses. (a) Experimental design for assessing transcriptional responses to IFN-γ and other cytokines. (b) Scatter plots comparing gene expression between unstimulated and IFN-γ-, TNF-α-, IL-17A-, IL-13-, or IL-1β-stimulated samples at day 5 in keratinocytes and melanocytes. Selected genes are labeled. (c) Scatter plots comparing transcriptional responses to IFN-γ, IL-17A, and IL-13 between 2D-cultured keratinocytes (n = 3) and keratinocytes isolated from skin biopsies after *ex vivo* cytokine stimulation of the biopsies (n = 1). Blue and red dots indicate genes significantly downregulated and upregulated, respectively in cytokine-stimulated 2D-cultured keratinocytes (|log2 fold change| > 1, FDR < 0.05). The five differentially expressed genes with the largest absolute log2 fold changes in 2D-cultured keratinocytes are labeled. Pearson’s *r* and Spearman’s *ρ* are shown for each comparison.

**Supplementary Figure 4.** Comparison of skin structural cell responses to primary or secondary IFN-γ stimulation. Frozen skin biopsies from healthy donors were subjected to primary (1°) or secondary (2°) IFN-γ stimulation. Skin structural cell populations were subsequently isolated by cell sorting and subjected to bulk RNA-seq to assess similarities and differences in transcriptional responses between 1° and 2° stimulation. (a) Scatter plots comparing log2 fold change following 1° and 2° IFN-γ stimulation relative to unstimulated samples in the indicated cell types. Light blue and light orange dots indicate genes significantly differentially expressed only following 1° and 2° IFN-γ stimulation, respectively, whereas blue and orange dots indicate genes significantly downregulated and upregulated following both 1° and 2° IFN-γ stimulation, respectively. Gray dots indicate all other genes. Differential expression was defined as |log2 fold change| > 1, FDR < 0.05. Pearson’s *r* and Spearman’s *ρ* and liner regression slopes are indicated. Dashed lines indicate the identity line (x = y). (b) Volcano plots of differential gene expression in the indicated cell types comparing 2° with 1° IFN-γ-stimulated samples. Blue and orange dots indicate genes significantly downregulated and upregulated, respectively, in 2° stimulated samples. IFN-γ-stimulated samples. Up to five significant genes (log2 fold change| > 1, FDR < 0.05) with the lowest FDR are labeled for each cell type Dotted lines indicate the thresholds for differential expression (|log2 fold change| > 1, FDR < 0.05). RNA-seq was performed using samples from n = 3 independent donors per cell type and condition.

**Supplementary Figure 5.** Keratinocyte enrichment and quality control of bulk ATAC-seq. Frozen skin biopsies from healthy donors were cultured *ex vivo* under the indicated IFN-γ stimulation conditions, and keratinocytes were subsequently enriched and subjected to bulk ATAC-seq. (a) Flow cytometry plots of keratinocyte before (top) and after (bottom) enrichment. ATAC-seq quality control (QC) metrics, including (b) fragment size distribution, (c) transcription start site (TSS) enrichment, (d) fraction of reads in peaks (FRiP) score, and (e) estimated library complexity. ATAC-seq was performed using samples from n = 3 independent donors per condition. Each symbol represents an individual donor.

