## Supplementary figures and images for "Establishing an *ex vivo* platform for molecular profiling of skin structural cells using frozen human skin biopsies"

### Fig S1

Supplementary Figure 1

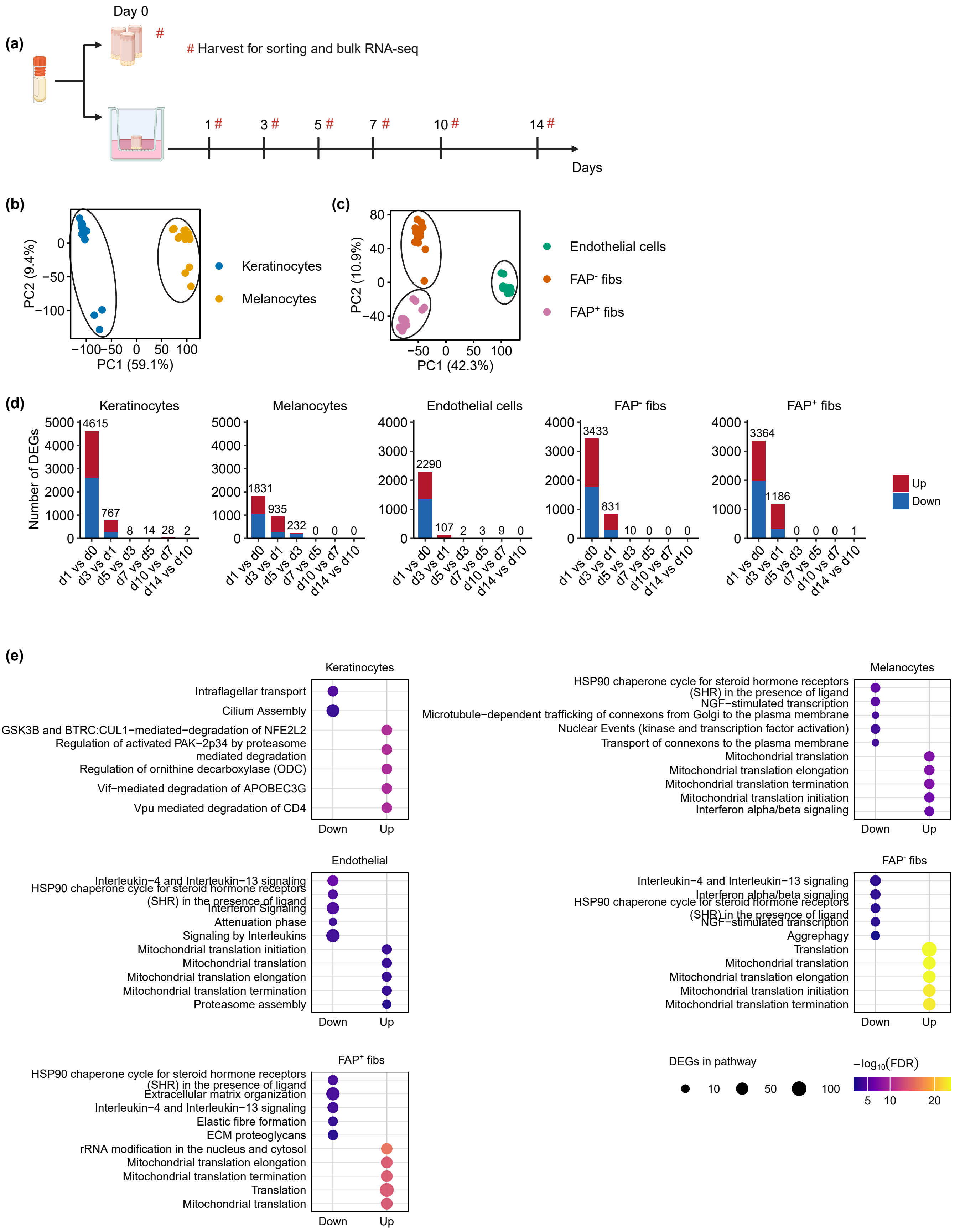

### Fig S2

#### Supplementary Figure 2

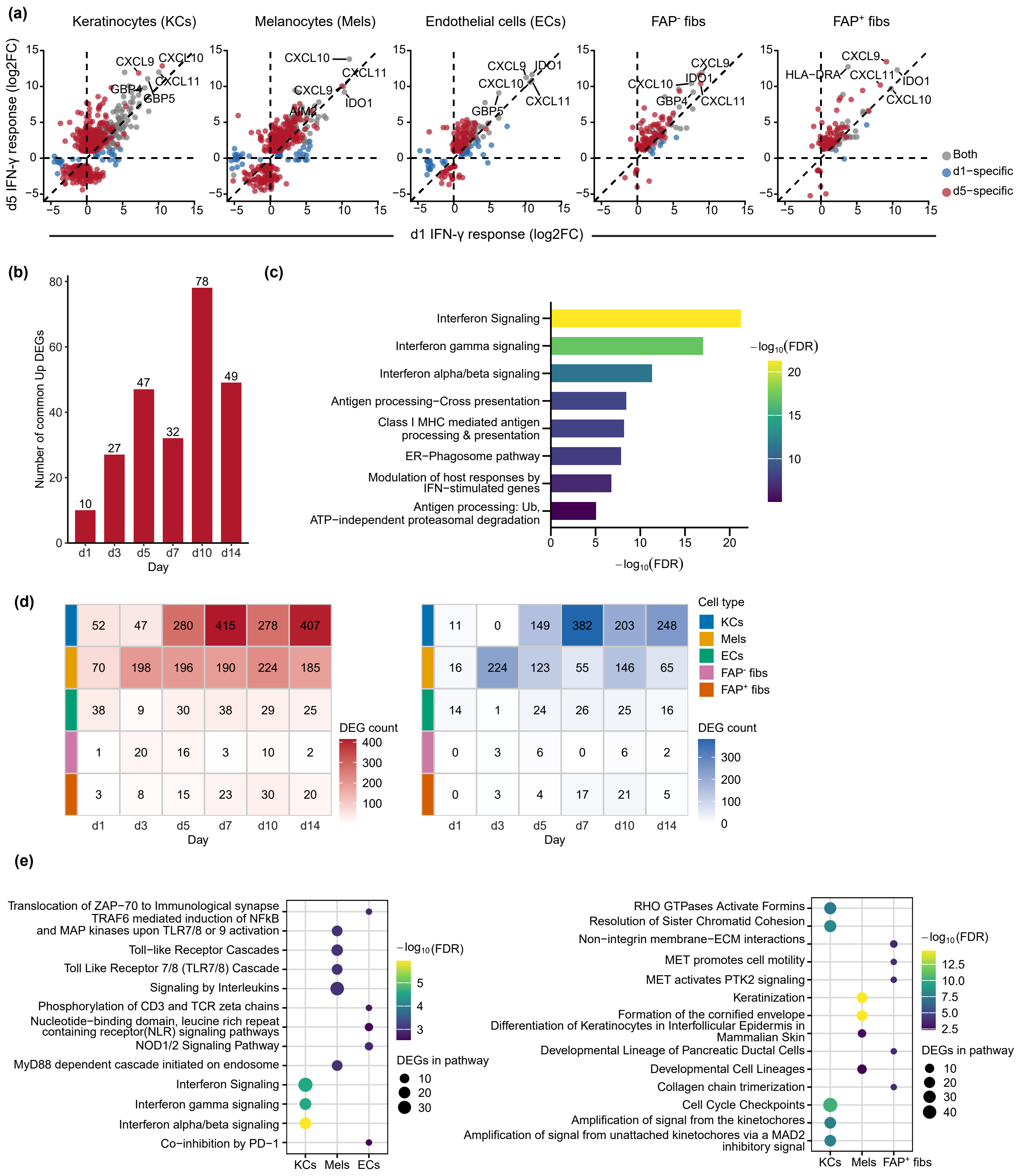

### Fig S3

Supplementary Figure 3

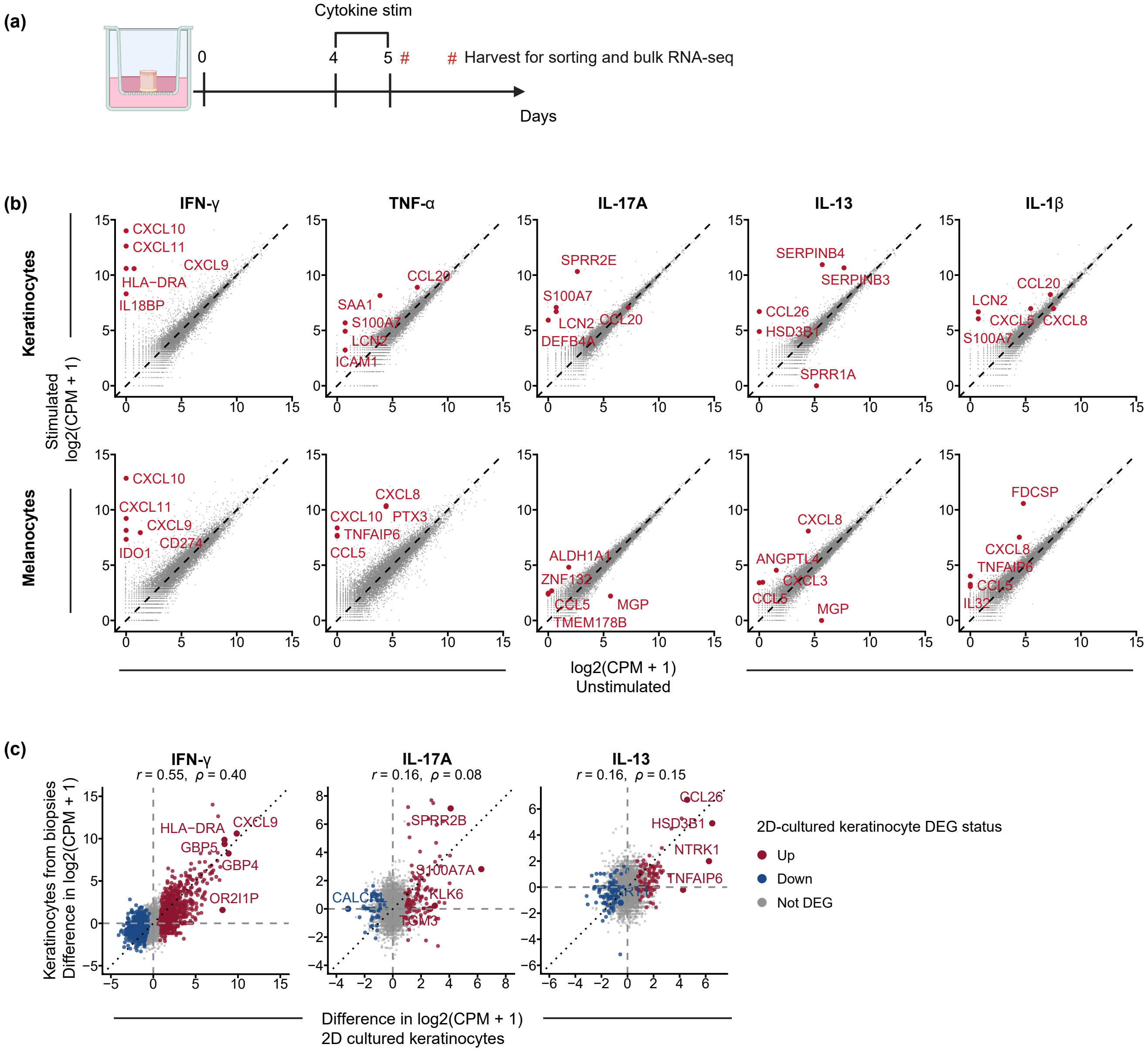

### Fig S4

Supplementary Figure 4

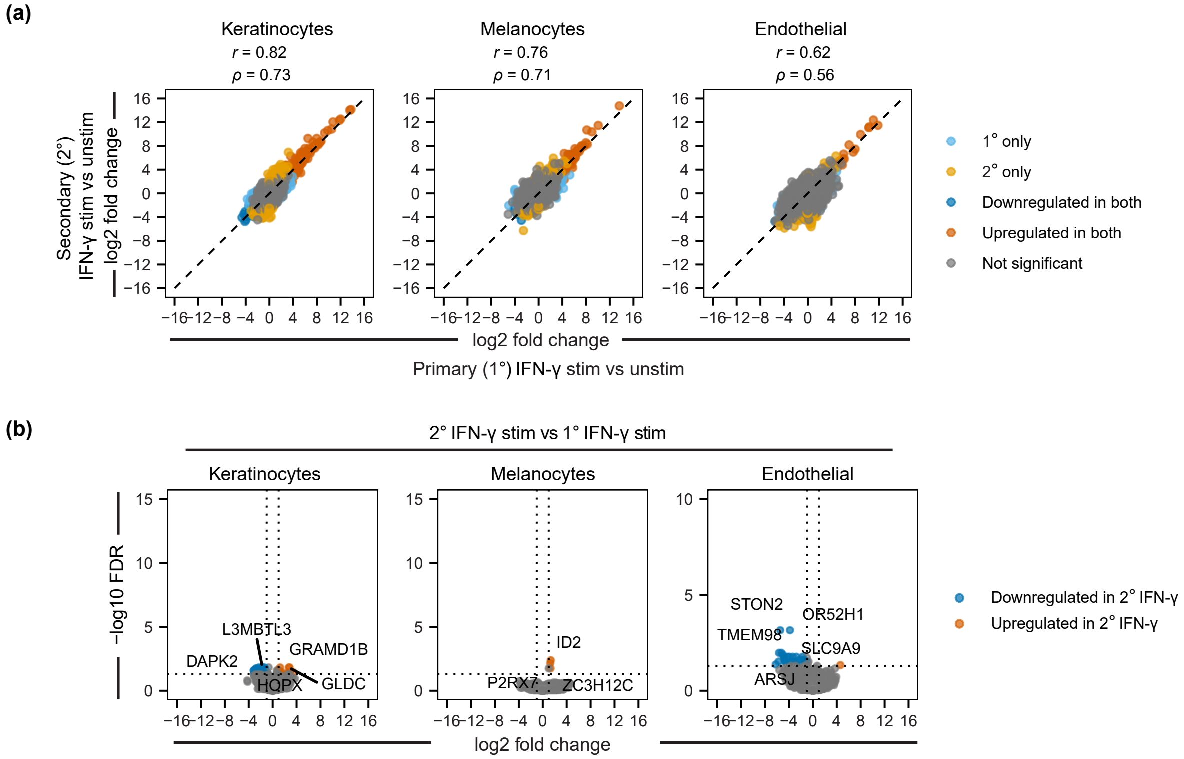

### Fig S5

Supplementary Figure 5

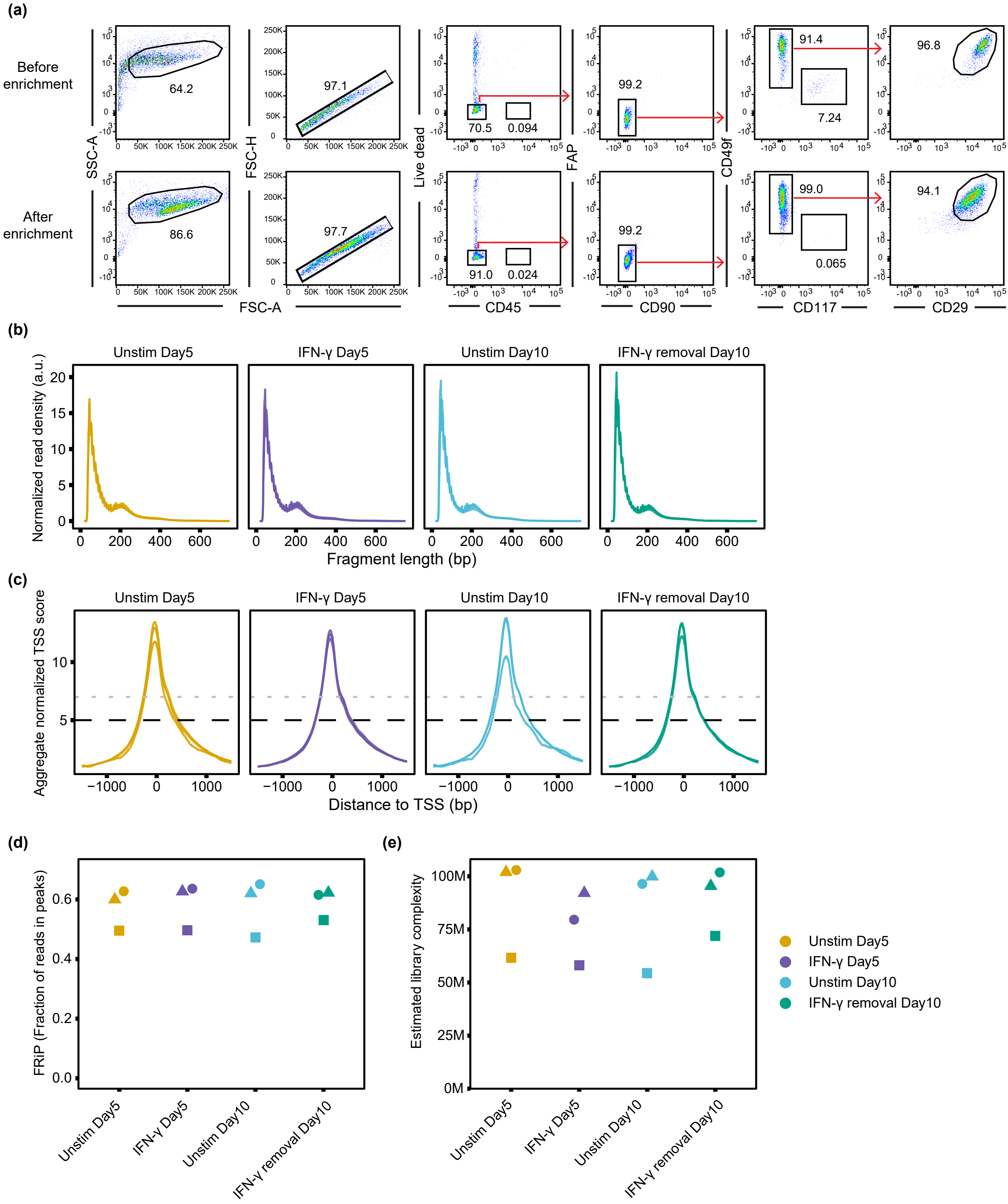
